# A novel method for large-scale identification of polymorphic microsatellites through comparative transcriptome analysis

**DOI:** 10.1101/114645

**Authors:** Wei Luo, Hongyue Qu, Xin Wang, Qin Zhan, Qiang Lin

**Affiliations:** CAS Key Laboratory of Tropical Marine Bio-resources and Ecology, South China Sea Institute of Oceanology, Chinese Academy of Sciences, Guangzhou, 510301, China.; University of Chinese Academy of Sciences, Beijing, 100049, China.; State Key Laboratory of Tropical Oceanography, South China Sea Institute of Oceanology, Chinese Academy of Sciences, Guangzhou, 510301, China.

**Keywords:** Micro satellites (SSR), Polymorphic, Transcriptome, Comparative transcriptome analysis, Sequence alignment

## Abstract

Microsatellite (SSR) is one of the most popular markers for applied genetic research, but generally the current methods to develop SSRs are relatively time-consuming and expensive. Although high-throughput sequencing (HTS) approach has become a practical and relatively inexpensive option so far, only a small percentage of SSR markers turn out to be polymorphic. Here, we designed a new method to enrich polymorphic SSRs through the comparative transcriptome analysis. This program contains five main steps: 1) transcriptome data downloading or RNA-seq; 2) sequence assembly; 3) SSR mining and enrichment of sequences containing SSRs; 4) sequence alignment; 5) enrichment of sequences containing polymorphic SSRs. A validation experiment was performed and the results showed almost all markers (> 90%) that were indicated as putatively polymorphic by this method were indeed polymorphic. The frequency of polymorphic SSRs was significantly higher (*P* < 0.05) but the cost and running time were much lower than those of traditional and HTS approaches. The method has a practical value for polymorphic SSRs development and might be widely used for genetic analyses in any species.

## INTRODUCTION

Microsatellites (SSRs) have been emerged as one of the most popular markers for a wide range of applications in population genetics, conservation biology and marker-assisted selection (Abdelkrim et al., 2009; Luo et al., 2012). Classically, microsatellite marker development requires: the construction of a genomic library enriched for repeated motifs; isolation and sequencing of microsatellite containing clones; primer design; optimization of PCR amplification for each primer pair; and a test of polymorphism on a few unrelated individuals. Most of these steps are either expensive, time-consuming, or both. With the wide application of high-throughput sequencing (HTS) technology, especially the whole transcriptome sequencing, development SSRs by HTS has become a practicable alternative for many species in recent years (Wu et al., 2014). It has greatly reduced the running time and cost requirement for SSR development. However, the frequency of polymorphic SSR markers developed by this method is much low in some species, which means that most of the loci cannot be effectively applied in genetic analysis (Iorizzo et al., 2011; Luo et al., 2012). According to our best knowledge, there is no records addressed the low frequency of polymorphic SSRs. Here we provided a new method for development of polymorphic SSRs through comparative transcriptome analysis.

## RESULTS AND DISCUSSION

Three, two and four transcriptomes of rice, grass carp and lined seahorse respectively were used and a total of 299, 206 and 956 putatively polymorphic SSRs were obtained by this method, respectively (Table 1; Table S1). Twenty, thirty and sixty loci were randomly selected for primer design, and 19 (95.00%), 26 (92.86%) and 50 (90.91%) loci showed polymorphic in rice, grass carp and lined seahorse, respectively (Table 1; Table S2). One-way ANOVA showed the frequency of polymorphic SSRs identified by this method was significantly (*P* < 0.05) higher than that of traditional approach and HTS (Fig. 2). In addition, we recently developed polymorphic SSR markers for lined seahorse by HTS approach, and the ratio of polymorphic SSRs was 17.93% (Arias et al., 2016). While using the same transcriptomes to develop SSRs by this method, the ratio was raised to 90.91%.

**Table 1.** Number of SSRs, polymorphic SSRs frequency of three verified species.

|  | Species for verification |  |  |
| --- | --- | --- | --- |
|  | Rice | Grass carp | Lined seahorse |
| Source of the transcriptome | SRR1799209<br>SRR1974265<br>SRR2048540 | SRR1618540<br>SRR1618542 | SCSIO-CAS |
| Total number of SSRs in transcriptome | 29,517 | 21,959 | 19,006 |
| Number of putatively polymorphic SSRs enriched by this program | 299 | 206 | 600 |
| Number of primers designed for validation experiments | 20 | 30 | 60 |
| Number of primers amplified with clear product bands (%) | 20 (100%) | 28 (93.33%) | 55 (91.7%) |
| Number of polymorphic SSRs (%) | 19 (95.00%) | 26 (92.86%) | 50 (90.91%) |

This method, which is based on the idea of enriching homologous sequences containing the same SSR with a different number of repeats, could identify polymorphic SSRs directly through comparative transcriptome analysis. Compared with traditional methods and HTS, this method have eliminated the most intensive wet lab steps, and the time and cost for primer synthesis and experimental validation by identifying and separating the many “monomorphic” SSRs from the minority polymorphic ones, cutting the running time and cost by half or more (Tang et al., 2008; Iorizzo et al., 2011). The fact that almost all tested SSRs predicted to be polymorphic were indeed validated as polymorphic demonstrates that it is an efficient and reliable method to develop polymorphic SSR markers. The method will play an important role in developing polymorphic SSR markers, providing a better service for the selective breeding and genetic studies.

## MATERIALS AND METHODS

### Materials

Each ten specimens of rice (*Oryza sativa*), grass carp (*Ctenopharyngodon idella*) and lined seahorse (*Hippocampus erectus*) were used to investigate the ratio of polymorphic SSRs developed by our method. The leaves of rice and the dorsal fin of grass crap and seahorse were used for DNA extraction.

### Architectural structure

The pipeline of this method consists of five steps (Fig. 1): 1) transcriptome data downloading or RNA-seq; 2) sequence assembly; 3) SSR mining and enrichment of SSR containing sequences; 4) sequence alignment; 5) enrichment containing polymorphic microsatellite sequences.

**Figure 1.**
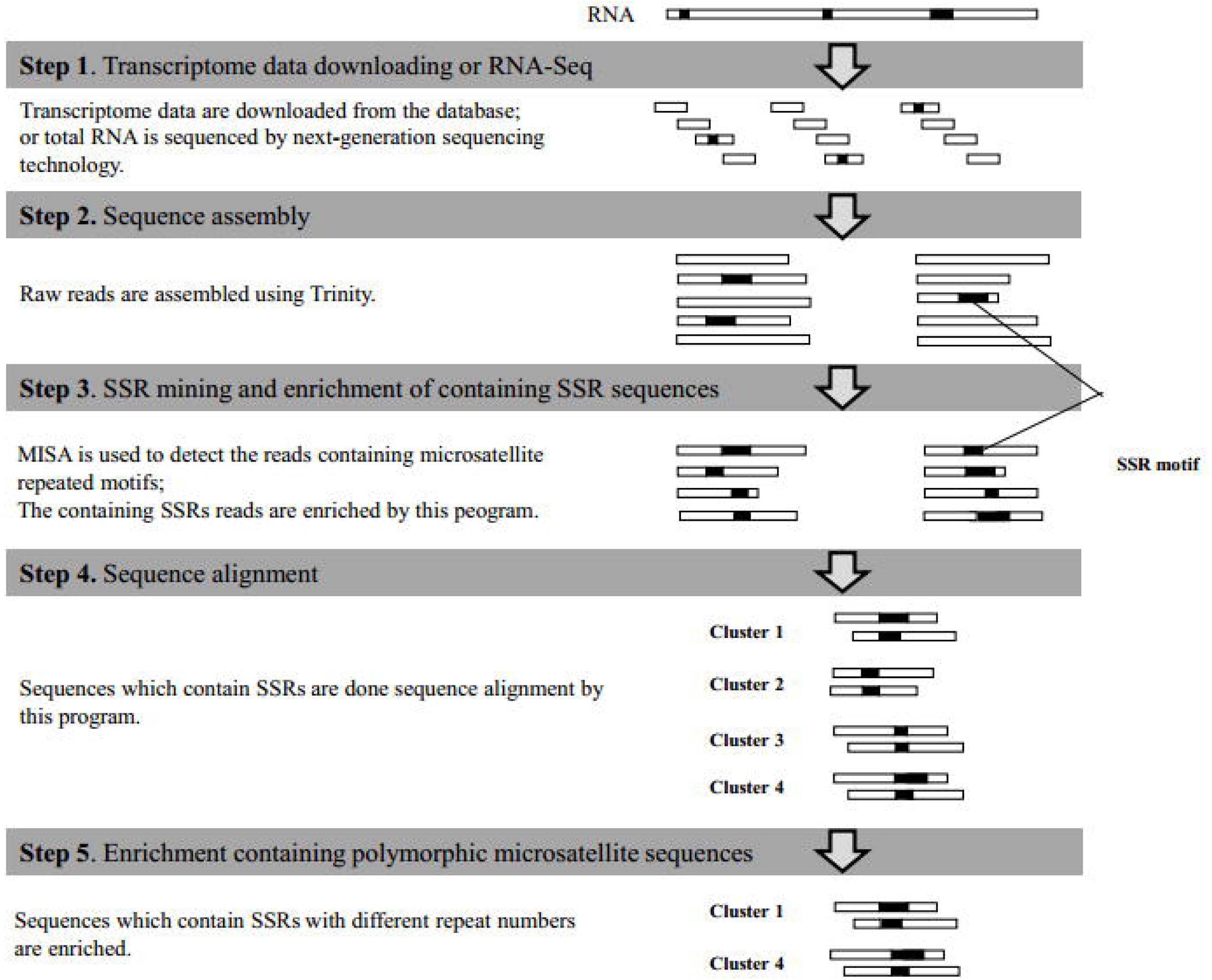
Flowchart of polymorphic markers enrichment and development.

**Figure 2.**
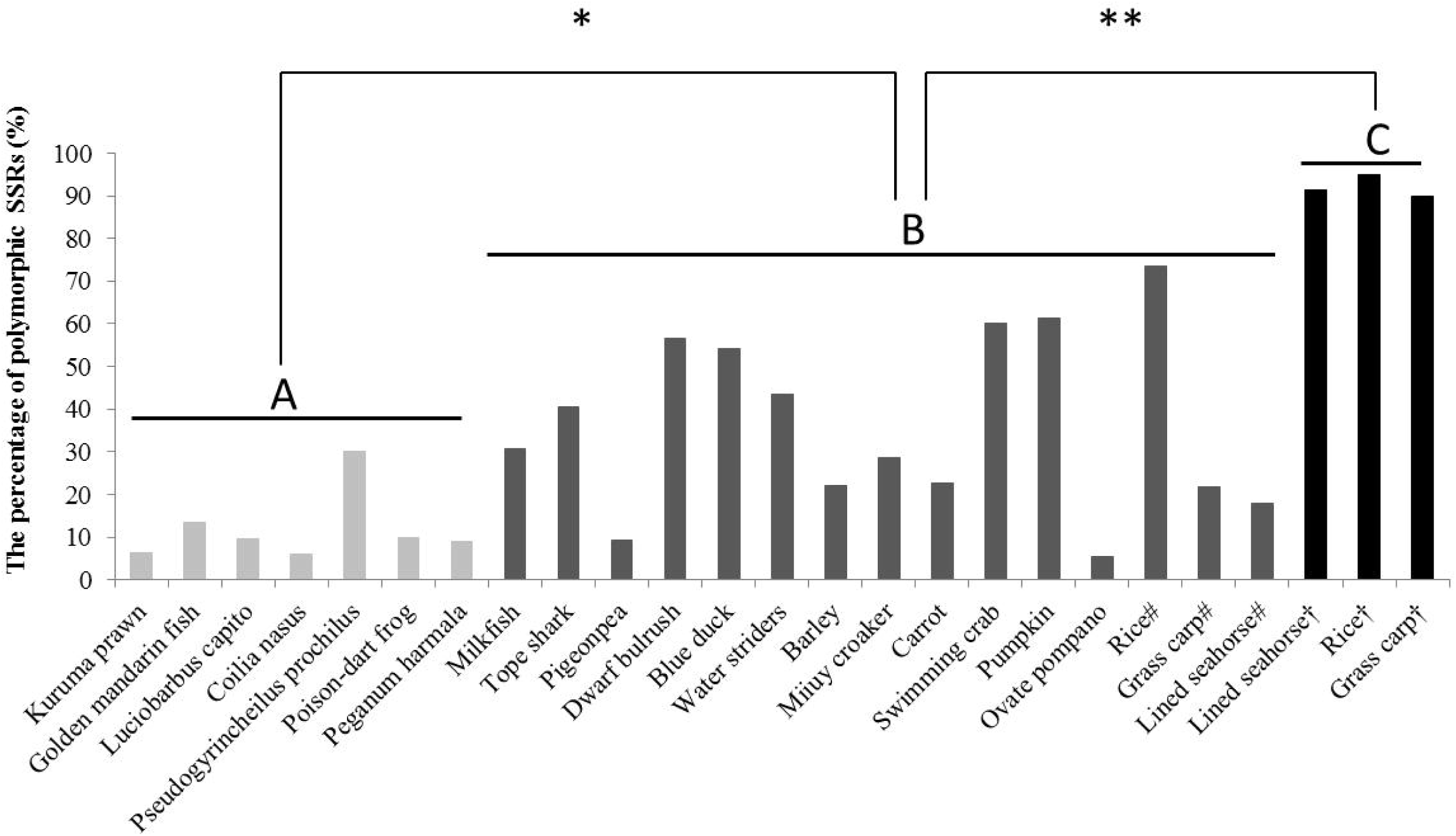
Comparison of polymorphic marker frequency developed by different methods. A, B and C represents SSRs developed by traditional methods, HTS approach and the method designed in this study, respectively; * and ** represents significance at *P* = 0.05 and *P* = 0.01, respectively; # represents SSR developed by HTS; † represents SSR developed by this method. The data of the frequency of polymorphic SSRs used in the figure was cited from the published references (Table S1)

### Transcriptome data gaining

The precondition of this method is to detect polymorphic SSRs in two or more transcriptomes from different samples. Three and two transcriptomes of rice and grass carp were downloaded from NCBI (Table 1). Four transcriptomes of seahorse were sequenced by us.

### De novo assembly

The raw reads were trimmed and quality controlled by SeqPrep (https://github.com/jstjohn/SeqPrep). Then clean data was used to perform RNA *de novo* assembly with Trinity using default parameters.

### SSR mining and enrichment of SSR containing sequences

We took rice as an example to enrich polymorphic SSRs. The three transcriptomes assembled were renamed “T1”, “T2” and “T3”, respectively.

MIcroSAtellite identification tool (MISA; http://pgrc.ipk-gatersleben.de/misa) was employed for SSR mining from different transcriptomes with the following settings (SSR motifs and number of repeats): dimer-6, trimer-5, tetramer-5, pentamer-5 and hexamer-5. In order to reduce the rate of false positives, a Python code was written to rule out the sequences which only contain mononucleotide repeats, compound SSRs, or end with SSRs. The command line is written as follows:

~~~
     *from Bio import SeqIO*
     *import os*
     *samples=['T1','T3','T4']*
     *for sample in samples:*
     *ids=[]*
     *faD=SeqIO. to_dict(SeqIO.parse (open(sample + '.fa'),'fasta'))*
     *for la in open(sample + '.ssr'):*
          *if'ID' not in la:*
               *aL=la.split('\t')*
               *ln=len (faD[aL[0]]. seq)*
               *if aL[2]!='p1' and aL[5]!=1 and aL[6]!=ln:*
                    *ids.append (aL[0])*
     *comp_ids=[]*
     *for lb in open(sample + '.ssr'):*
          *bL=lb. strip (). split ('\t')*
          *if'c' in bL[2]:*
               *comp _ids.append (bL[0])*
     *fas=SeqIO.parse(open(sample + '.fa'), fasta*)
     *for fa in fas:*
          *if fa.id in ids and fa.id not in comp_ids:*
               *open(sample + '.ssr.fa','a').write('>%s\n%s\n' % (fa.id,str(fa.seq)))*
~~~

And then we renamed all the transcriptomes and all the sequences containing SSRs detected with MISA software by adding different prefixes. Finally, we combined all the sequences containing SSRs from different transcriptomes into a file. The command line is written as follows:

~~~
          *from Bio import SeqIO*
          *samples=['T1','T3','T4]*
          *for sample in samples:*
          *fas=SeqIO.parse(open(sample+'.Trinity.fasta'), 'fasta')*
          *for fa in fas:*
               *open(sample+'.fa','a').write(>%s.%s\n%s\n'%(sample,fa.id,str(fa.seq)))*
          *for la in open(sample+'.Trinity.fasta.misa'):*
               *if'ID' not in la:*
                    *la =sample+'.'+la*
          *open(sample + '.ssr', 'a').write(str(la))*
~~~

### Alignment of containing SSR sequences

Sequences containing SSRs were clustered using the default parameters of the CD-HIT tool at 90% sequence identity level.

### Rename the SSR file

We then edited a Python code that generated the reverse complement of the minus strand transcripts, according to the strand information in the output of CD-HIT:

~~~
     *import re*
     *from Bio import SeqIO*
     *faD=SeqIO.to_dict(SeqIO.parse(open('all.ssr.fa'), 'fasta'))*
     *baseD={'A ':'T', 'T':'A', 'G':'C', 'C':'G'}*
     *samples=['T1','T3','T4']*
     *for sample in samples:*
     *for line in open(sample+'.ssr'):*
          *lst=line. strip ().split('\t')*
          *if 1st[0] in faD:*
               *if 'c' not in lst[2] and lst[2]! = 'p1' and 'ID' not in line and lst[-2]!="1" and int(lst[-1])!=len(faD[lst[0]].seq) and int(lst[-1])!=len(faD[lst[0]].seq)-1 and int(lst[-1])!=len(faD[lst[0]].seq)-2 and int(lst[-1])!=len(faD[lst[0]].seq)-3:*
                    *ma=re.findall('\((.+)\)', lst[3])[0]*
                    *maL=list(ma)*
                    *rc = "*
                    *for base in maL:*
                         *if base in 'ACGT':*
                              *rc + =baseD[base]*
                    *rc=rc[::-1]*
                    *ss=[ma,rc]*
                    *ss.sort()*
                    *ss='('+ '/'.join(ss)+*)'
                    *newline =re.sub('\(([ACGT] +)\) ',ss,line)*
                    *open(sample+'.ssr.reformed', 'a').write(str(newline))*
~~~

And then, we edited a Python code that generated reverse complementary of minus strand transcripts, according to the strand information in the ouput of CD-HIT:

~~~
     *from Bio import SeqIO*
     *import re*
     *sD={}*
     *for la in open('cd-hit.clstr'):*
     *if'at' in la:*
          *id=re.findall('> (+)\. \.\.',la)[0]*
          *strand=re.findall ('([+-]) + V',la)[0]*
          *sD[id]=strand*
     *if'*' in la:*
          *id=re.findall('> (.+)\. \.\.',la)[0]*
          *sD[id]='+'*
     *fas =SeqIO.parse(open('all.ssr.fa'), fasta')*
     *for fa in fas:*
     *if fa.id in sD:*
          *if sD[fa.id]= = '+':*
               *open('plus.ssr.fa', 'a').write('>'+str(fa.id) + '\n'+str(fa.seq) + '\n')*
          *if sD[fa.id]= = '-':*
               *seq=fa.seq.reverse_complement()*
               *open('plus.ssr.fa', 'a').write('>'+str(fa.id)+'\n'+str(seq)+'\n')*
~~~

### Enrichment of sequences containing polymorphic SSRs

A script was executed to enrich sequences with different repeats from all the sequences containing SSRs:

~~~
     *import re*
     *from collections import defaultdict*
     *from Bio import SeqIO*
     *import os*
     *def getD(ssr):*
     *s=[]*
     *for la in open(ssr):*
          *if'ID' not in la:*
               *aL=la.strip (). split('\t')*
               *ma=re.findall('\(.+\)\d+',aL[3])*
               *s.append ((aL[0],ma[0]))*
     *d=defaultdict(set)*
     *for k, v in s:*
          *d[k].add(v)*
     *return d*
     *t1D=getD('T1.ssr.reformed')*
     *t2D =getD ('T3.ssr. reformed')*
     *t3D=getD('T4.ssr.reformed')*
     *allD={}*
     *allD.update(t1D)*
     *allD.update(t2D)*
     *allD.update(t3D)*
     *page =open ('cd-hit. clstr').read ()*
     *clusters=re.findall ('(. + ?)>Cluster ',page,re.S)*
     *for cluster in clusters:*
     *trans=re.Jindall('T\d\.TR\d+\\c\d+_g\d+_i\d+',cluster,re.S)
     if len(trans)>1:*
          *tt=[]*
          *ss=[]*
          *for tran in trans:*
               *if tran in allD:*
                    *tt.append(str(tran) +':'+str(list(allD[tran])))*
                    *ss+=list(allD[tran])*
          *if len(tt)>1:*
               *ma=re.findall('\)\d+\",str(ss))*
               *ma=set(ma)*
               *mas=re.findall('\((. + ?)\)',str(set(ss)))*
               *ssr=”*
               *for mm in list(set(mas)):*
                    *if mas.count (mm)> 1:*
                         *ssr=mm*
               *if len(ma)>1 and len(mas)>len(set(mas)):*
                    *ttt=[]*
                    *for t in tt:*
                         *if ssr in t:*
                              *ttt.append(t)*
                    *na=str(ttt).lstrip('[').rstrip(']').strip("").replace("', "', '\t)*
                    *open('enrichment.SSRs','a').write(str(na)+'\n')*
     *for 11 in open('enrichment.SSRs'):*
     *ma=re.findall('([a-zA-Z0-9]+)\.TR\d+',11)*
     *ma=set(ma)*
     *iflen(ma)>1:*
          *open('enrichment.SSRs.txt','a').write(str(11)) faD=SeqIO.to_dict(SeqIO.parse(open('plus.ssr.fa'), 'fasta'))*
     *n=0*
     *for la in open('enrichment.SSRs.txt'):*
     *aL=la. strip (). split ('\t')*
     *for it in aL:*
          *id=it. split (': ')[0]*
          *open('cluster'+str(n), 'a').write('>'+str(id)+'\n+strfaD[id].seq)+'\n*)
     *os.system('muscle -msf -in cluster'+str(n)+' -out cluster'+str(n) + '.muscle')*
     *n+=1*
*os.system('mkdir muscle; mv cluster* muscle; rm enrichment.SSRs')*
~~~

## Validation experiments

Primers were designed using Primer premier 5 software. The PCR products were separated by capillary gel electrophoresis using the ABI 3100 Genetic Analyser. The peak heights and fragment sizes were analyzed using GeneMarker software.

## Data analysis

Previous studies on SSR development in animals and plants, along with their frequency of polymorphic SSR markers, were randomly downloaded from the internet (Table S3). Differences in mean value of the frequency of polymorphic SSRs developed by three methods were analyzed using one-way analysis of variance (ANOVA).

## Competing interests

The authors declare no conflict of interest.

## Author contributions

W.L. and Q.L. conceived and designed the experiments; H.Q., X.W. and Q.Z. performed the experiments and analyzed data; W.L., and Q.L. prepared the manuscript. All authors read and approved the final manuscript.

## Funding

This study was funded by the National Natural Science Foundation of China (No. 41576145) and the Guangdong Oceanic and Fisheries Science and Technology Foundation (No. A201501A12).

## Supplementary information

**Table S1.** The putative polymorphic SSRs enriched by this method in rice (*O. sativa*), grass carp (*C. idella*) and lined seahorse (*H. erectus*).

**Table S2.** The SSR loci validated by experiments and characteristics of the primers.

**Table S3.** The species and the frequency of polymorphic SSRs cited in this study.

